# Conversion of Genomic DNA to Proxy Constructs Suitable for Accurate Nanopore Sequencing

**DOI:** 10.1101/027284

**Authors:** Dimitra Tsavachidou

## Abstract

Nanopore sequencing at single-base resolution is challenging. There are developing technologies to convert DNA molecules to expanded constructs. Such constructs can be sequenced by nanopores in place of the original DNA molecules. We present a novel method for converting genomic DNA to expanded constructs (“proxies”) with 99.67% accuracy. Our method “reads” each base in each DNA fragment and appends an oligonucleotide to the DNA fragment after each base “reading”. Each appended oligonucleotide represents a specific base type, so that the proxy construct consisting of all the appended oligonucleotides faithfully represents the original DNA sequence. We generated proxies for genomic DNA and confirmed the identities of both the proxies and their corresponding original DNA sequences by performing sequencing using Ion Torrent sequencer.

Conversion to proxies had only 0.33% raw error rate. Errors were: 93.96% deletions, 5.29% insertions, and 0.74% substitutions. The longest sequenced proxy was 170 bases, corresponding to a 17-base original DNA sequence. The short length of the detected proxies reflected restrictions imposed by Ion Torrent’s short reads and was not caused by limitations of our method. The consensus sequence built by using proxies alone (average length: 120 bases; corresponding to original sequences with average length 12 bases) covered 55% of the reference genome with 100% accuracy, and outperformed the Ion Torrent sequencing of the corresponding original DNA fragments in terms of accuracy, coverage and number of aligned sequences. Data and other materials can be found at http://www.vastogen.com/data.html. This proof-of-concept experiment demonstrates highly accurate proxy construction at the whole genome level. To our knowledge, this is the first demonstrated construction of expanded versions of DNA at the whole genome level.

## Introduction

A promising technology that has the potential to revolutionize sequencing by simplifying the process and lowering the cost is nanopore-based detection. Nanopore devices are able to differentiate between short DNA segments with distinct sequences, but they have difficulty performing sequencing at single-nucleotide resolution. Sequencing at single-nucleotide resolution is not feasible with solid-state nanopores, and is performed with high error rates when using biological nanopores (Goodwin et al., 2015).

Problems arising from nanopore sequencing at single-nucleotide resolution can be circumvented by using expanded versions of nucleic acid molecules that can be readily detected by nanopore devices. Such expanded constructs preserve the sequence information of the nucleic acid molecules that they represent. Methods to generate expanded versions of nucleic acid molecules have been proposed previously, but they are difficult to use, because they are based on inefficient circularization steps (Lexow, 2008)(Buzby et al., 2012)(Meller and Weng, 2012), or on complex and inefficient hybridization steps, or expandable nucleotides with complex structures (Kokoris and McRuer, 2013).

Here we present results using a novel method to construct expanded versions of genomic DNA molecules called “proxies”.

## Results and Discussion

### Proxy construction

A proxy is an expanded version of a DNA molecule that faithfully represents the DNA molecule’s sequence and consists of tail-tags (Figure 1A). In this study, proxies were constructed by using a novel method that (i) “reads” a first base close to one end of a DNA fragment, (ii) appends an oligonucleotide (“tail-tag”) (Figure 1B) to the other end of the DNA fragment, and (iii) repeats (i) and (ii) for the next base. Each tail-tag has a sequence that uniquely represents one of the four base types (A, T, C or G). For example, in the event that the base being “read” in step (i) is “A”, the appended tail-tag has a specific sequence that represents “A” uniquely.

**Figure 1.**
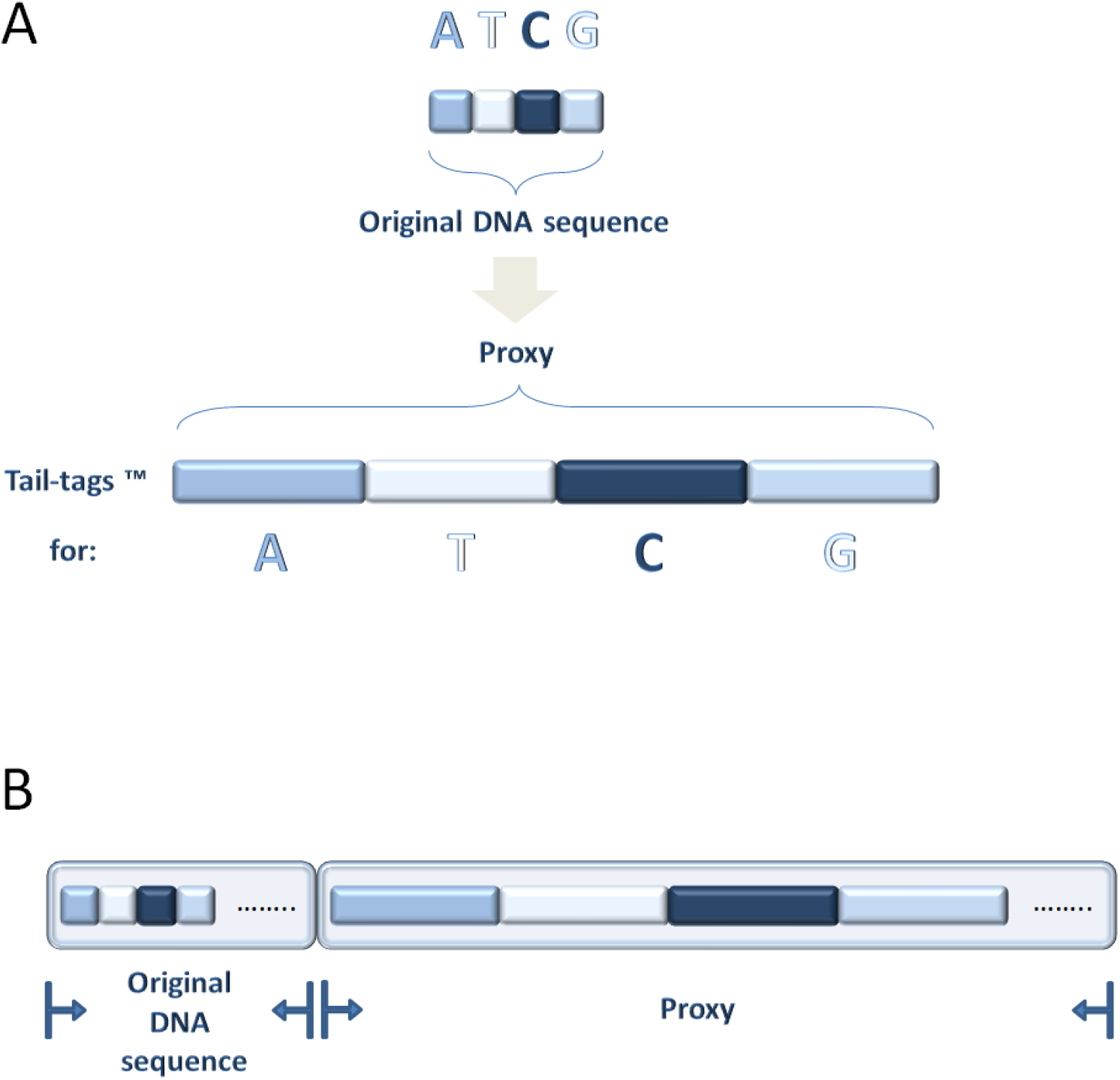
A: An original DNA sequence “ATCG” is shown converted to a proxy which consists of tail-tags, each of which uniquely represents A, T, C, or G. B: A proxy is shown appended to an original DNA sequence. Dots represent bases and tail-tags not shown in the drawing.

In this study, we conducted a proof-of-concept experiment, using lambda phage genomic material. The workflow is shown in Figure 2. Briefly, lambda phage genomic DNA was fragmented and attached to adaptors immobilized to magnetic beads. The immobilized adaptors comprised primer sequences suitable for Ion Torrent sequencing. Then, proxy construction was performed, followed by attaching adaptors comprising sequences compatible with immobilization to Ion Torrent emulsion PCR beads. Proxy construction was preceded by the addition of a separator (i.e. an oligonucleotide comprising a distinct sequence that marks the end of each genomic fragment and the beginning of its appended proxy).

**Figure 2.**
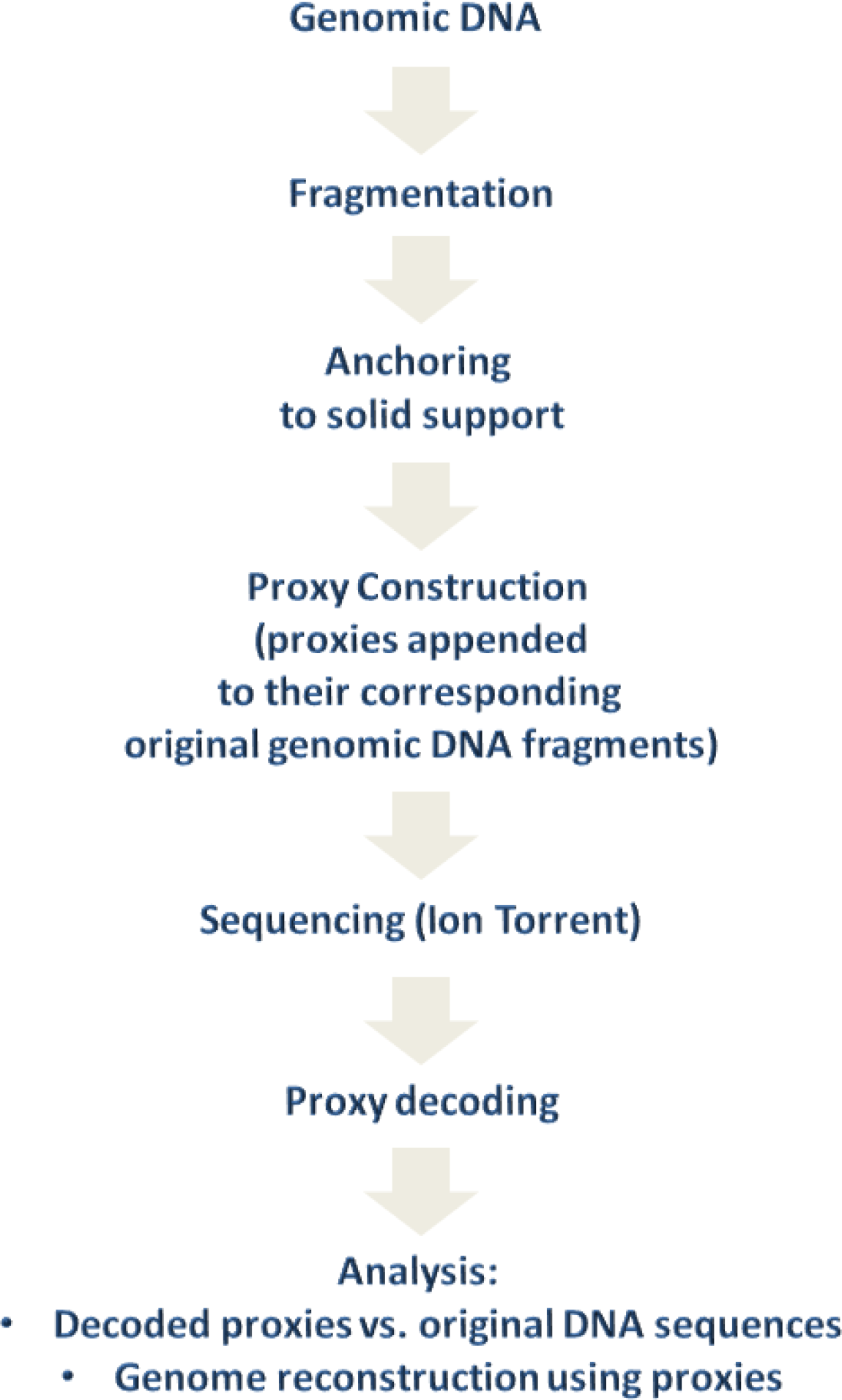
Experimental workflow

The principle of proxy construction is shown in Figure 3.

**Figure 3.**
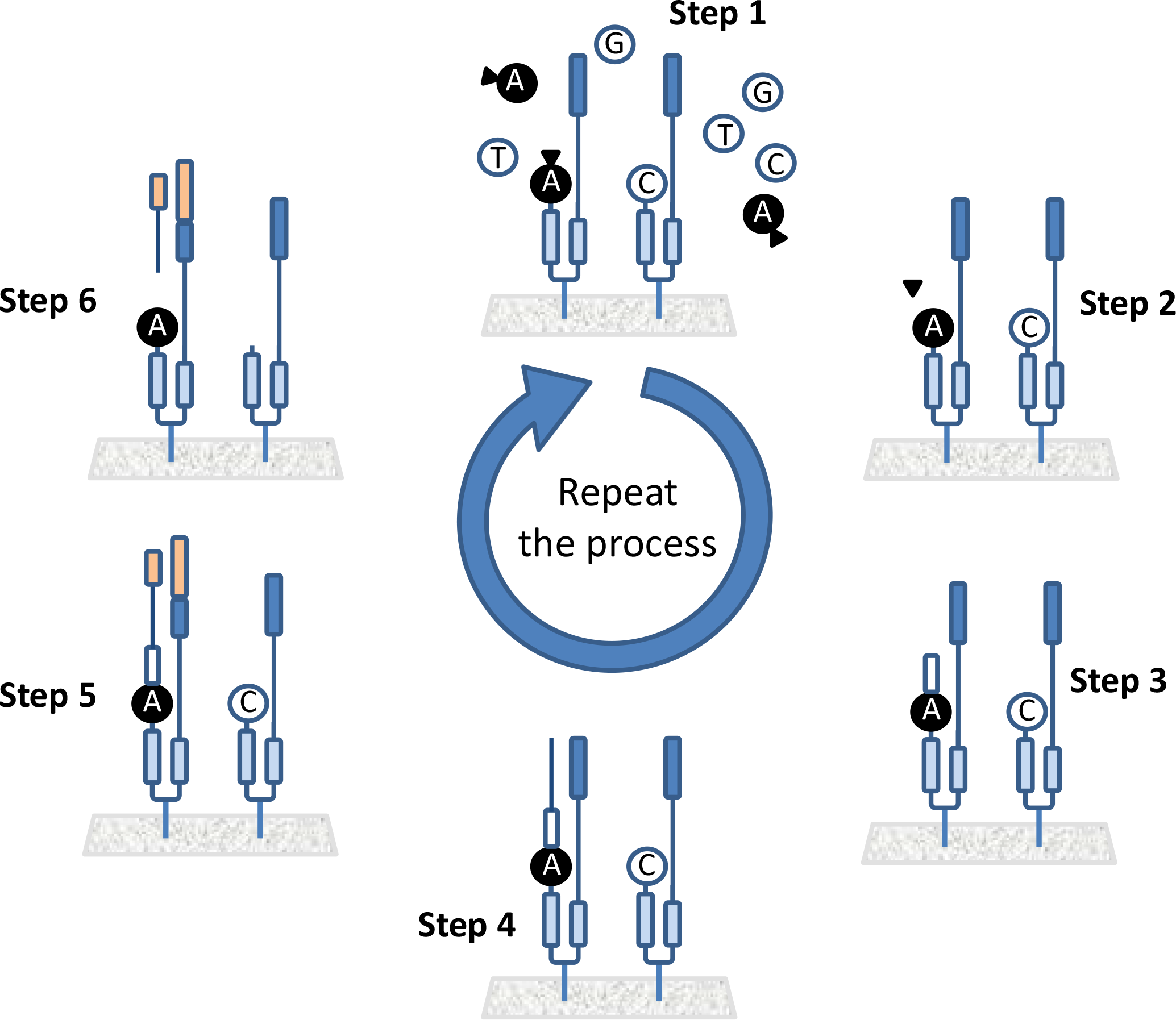
DNA proxy construction

During step 1, DNA extension is performed, using reversibly terminated nucleotides with only one base (e.g., “A”), and irreversibly blocked cleavable nucleotides with the other three bases.

During step 2, the reversible terminators of the reversibly terminated nucleotides are removed.

During step 3, removable extensions containing cleavable nucleotides are formed at extendable 3’ ends generated after removing the reversible terminators in the previous step.

Step 4 ensures that removable extensions in the previous step are fully extended.

During step 5, fully extended constructs from the previous step are properly cut and ligated to tail tags that correspond to the base type of the reversibly terminated nucleotides incorporated in step 1.

During step 6, any removable extensions formed during step 3, and any cleavable nucleotides incorporated during step 1 are removed, and the constructs are ready to participate in a next cycle of steps 1 through 6, to create proxies.

### Proxies were constructed with high accuracy

For the purpose of determining the identities of both the DNA fragments and their associated proxies, DNA fragments and their appended proxies were amplified and subjected to high-throughput sequencing using an Ion Torrent sequencer. The general structure of full-length sequenced products (excluding adaptors and the key sequence TCAG) is shown in Figure 4. Overall, 325,880 sequences were produced. The sequences (starting with the key sequence) are provided in Supplemental File 1. Proxies with their tail-tags were identified and decoded using custom parser scripts. Decoding was simply the conversion of each tail-tag back to the single-letter DNA code, A, T, C or G. Of the 325,880 sequences, 147,469 contained proxies with at least one tail-tag. The average number of tail-tags within a proxy was 5.6. The proxy size distribution is shown in detail in Table 1. The decoded proxy sequences and the original DNA sequences (at least part of which) the proxies represent are given in Supplemental File 2. It is important to note that the apparent small number of tail-tags is not due to limitations of our proxy construction method, but is a reflection of the short-read nature of Ion Torrent sequencing.

**Figure 4.**
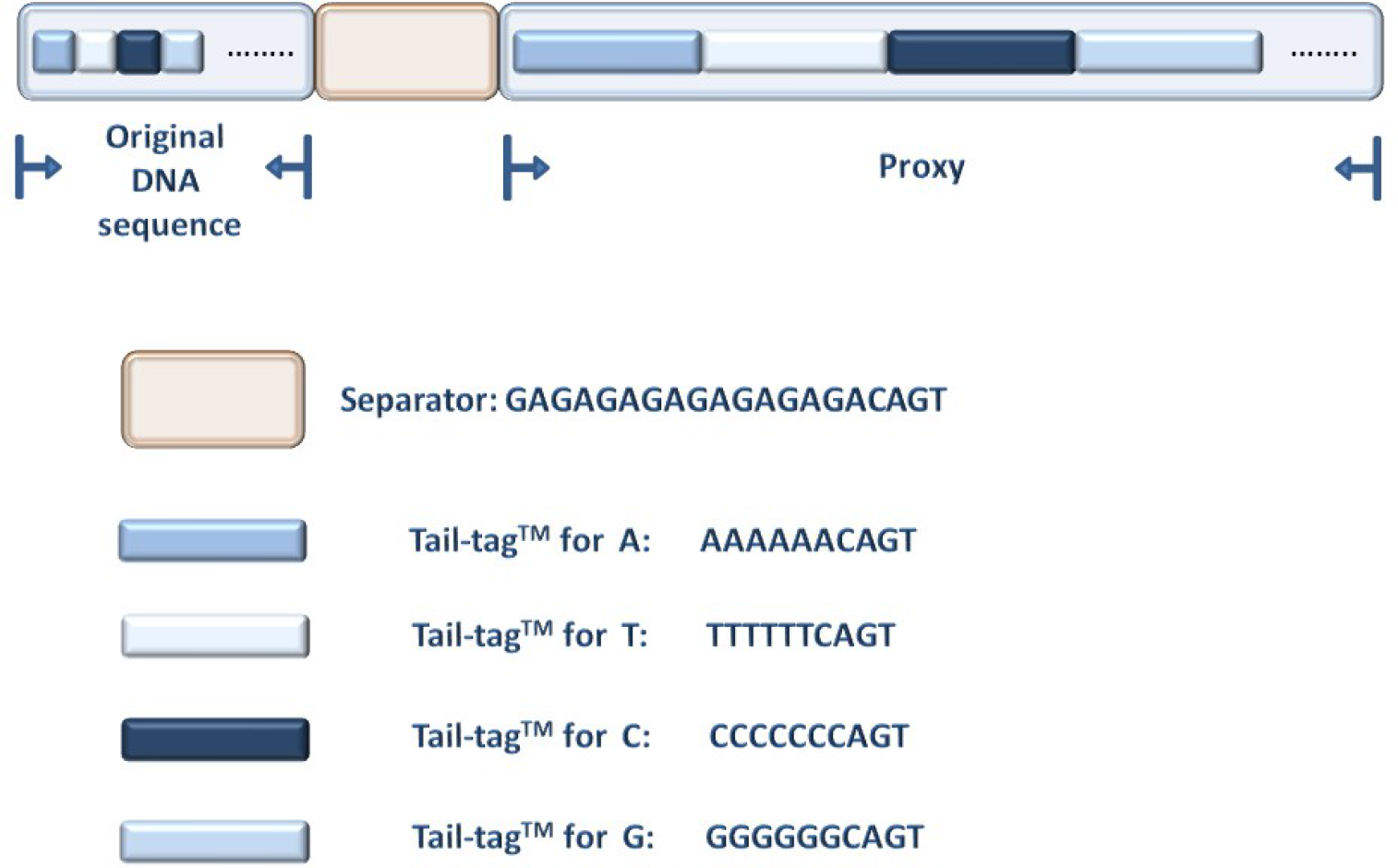
The general structure of an example full-length sequence is shown.

**Table 1.** The number of proxies (occurrence) of each examined length (number of tail-tags within the proxy) is shown.

| tail-tag number | occurrence |
| --- | --- |
| 1 | 32805 |
| 2 | 16193 |
| 3 | 12575 |
| 4 | 10763 |
| 5 | 9327 |
| 6 | 8945 |
| 7 | 9433 |
| 8 | 7786 |
| 9 | 6258 |
| 10 | 7114 |
| 11 | 7728 |
| 12 | 6517 |
| 13 | 4826 |
| 14 | 4808 |
| 15 | 2243 |
| 16 | 146 |
| 17 | 2 |

In order to assess how accurately the proxies represent their corresponding original DNA sequences, a comparison of decoded proxy sequences vs. original DNA sequences was performed using custom scripts.

It is important to note that the redundancy in the tail-tag and separator sequences allows for accurate decoding regardless of errors introduced by Ion Torrent sequencing. For example, seeking “GAGAGAGA” within a sequence locates the separator even if the separator was sequenced as “GAGAGAAGAGAGAGAGA” instead of “GAGAGAGAGAGAGAGA”. Similarly, looking for “AAA” located after the separator sequence identifies an “A” tail-tag when it is erroneously sequenced as “AAGAAA” instead of “AAAAAA”.

Direct comparison between original DNA sequences and decoded proxies revealed that approximately 95% of the proxies were error-free (i.e. correctly representing their corresponding original DNA molecules), and that the estimated raw error rate (the percentage of tail-tags not matching their corresponding bases in the original DNA sequence) was 0.97%. The number of correct proxies was underestimated and the raw error rate was inflated, because Ion Torrent introduced sequencing errors in the original DNA sequences that negatively affected the comparison between decoded proxies and original DNA sequences. In fact, Ion Torrent technology is reported to have 1.71% raw error rate (Quail et al., 2012).

In order to prevent Ion Torrent errors from affecting our results, we performed alignment of all the original DNA sequences to the reference genome using Bowtie. From the resulting SAM file, we determined which sequences were aligned, and how many bases from the starting position of each original DNA sequence perfectly matched the reference genome. We considered only the bases of the original DNA sequences that perfectly matched the reference genome, and compared them to their corresponding tail-tags. The original DNA sequences that aligned to the reference genome, the number of consecutively perfectly matched bases (starting from first base position) and the corresponding proxies are provided in Supplemental File 3.

Analysis revealed that 98.3% of the proxies were error-free (i.e. correctly representing their corresponding original DNA molecules), and that the estimated raw error rate (the percentage of tail-tags not matching their corresponding bases in the original DNA sequence) was only 0.33%. Further analysis showed that the errors were: 93.96% deletions, 5.29% insertions, and 0.74% substitutions.

### Genome reconstruction using proxies

In addition to comparing proxies to original DNA sequences, we also performed alignment of decoded proxies at least 10 bases long to the reference genome, as a different means of evaluating proxy performance. 33,384 decoded proxies were at least 10 bases long (average length: 12 bases), 98.8% of which aligned to the reference genome. The consensus sequence constructed by the aligned decoded proxies was determined by using Samtools and VarScan (to analyze the Bowtie-generated SAM file). The consensus sequence covered 55% of the reference genome with 100% accuracy.

Then, we aligned the sequences of the original DNA fragments; only the parts of the sequences that were represented by the above-described 33,384 proxies were used. A lower percentage of sequences (97.7%) aligned to the reference genome compared to the proxies, and the resulting consensus sequence covered 51.3% of the reference genome with 100% accuracy. The better performance of the proxies compared to the original DNA sequences reflects the advantage of the elongated nature of the proxies, which compensates for errors introduced during sequencing.

### Future steps and Conclusions

As mentioned previously in this manuscript, the short length of the decoded proxies is not the result of limitations of our method, but is a direct consequence of using Ion Torrent. Ion Torrent was chosen because of availability, low cost and ability to determine both the sequences of the proxies and their corresponding original DNA sequences. Future experiments will include constructing longer proxies, using longer and more tail-tags, and sequencing them using nanopores.

It is important to mention that the advantage that the proxies will confer to nanopore sequencing is not only higher accuracy rates, but also markedly improved speed. Current nanopore technologies perform sequencing base-by-base and do so by using complex chemistries to considerably slow down DNA passage through the pores. Proxies will bypass the need for complex chemistries and enable nanopore sequencing at native passage speeds, significantly increasing output.

To our knowledge, this is the first demonstrated accurate construction of expanded versions of DNA at the whole genome level. The very low raw error rate of proxy construction suggests that this method can become competitive in the field of genome sequencing.

## Methods and Supplemental Information

Supplemental Files 1-3 can be found at www.vastogen.com/data.html

Please contact the author at for questions or to request access to data files such as FASTQ and SFF.

## References

Buzby, P.R., Meller, A., Mcnally, B., Fan, A., Olejnik-Krzynmanska, E., 2012. Sequence preserved dna conversion for optical nanopore sequencing. US20120316075 A1.

Goodwin, S., Gurtowski, J., Ethe-Sayers, S., Deshpande, P., Schatz, M., McCombie, W.R., 2015. Oxford Nanopore Sequencing and de novo Assembly of a Eukaryotic Genome. bioRxiv 013490. doi:10.1101/013490

Kokoris, M.S., McRuer, R.N., 2013. United States Patent: 8349565 - High throughput nucleic acid sequencing by expansion. 8349565.

Lexow, P., 2008. Method for sequencing a polynucleotide. WO2008032058 A2.

Meller, A., Weng, Z., 2012. Sequence preserved dna conversion. US20120040869 A1.

Quail, M.A., Smith, M., Coupland, P., Otto, T.D., Harris, S.R., Connor, T.R., Bertoni, A., Swerdlow, H.P., Gu, Y., 2012. A tale of three next generation sequencing platforms: comparison of Ion Torrent, Pacific Biosciences and Illumina MiSeq sequencers. BMC Genomics 13, 341. doi:10.1186/1471-2164-13-341

